# Two peas in a pod: retroviral RNA dimers organize Gag-RNA nanoclusters with novel biophysical properties

**DOI:** 10.1101/2025.06.09.658653

**Authors:** Gregory S. Lambert, Christopher A. Siedlecki, Leslie J. Parent

## Abstract

The continued effective control of retroviral infection will no doubt require the development of new clinical interventions targeting underexploited areas of retroviral biology such as genome selection and virion assembly. In our previous work, we demonstrated that both the Gag-psi (Ψ) interaction and genomic RNA (gRNA) dimerization each uniquely contribute to the formation, morphology, and stability of RSV Gag-viral RNA (vRNA) biomolecular condensates (BMCs). The present work builds upon those observations, utilizing atomic force microscopy (AFM) and fluorescence correlation spectroscopy (FCS) to elucidate the nanoscale morphology, resistance to mechanical deformation, and constituent diffusivity of RSV Gag-vRNA BMCs. These approaches revealed a novel role for gRNA dimerization in nanoscale condensate architecture and mechanical stability that aids in our understanding of why gRNA dimerization is so critical for efficient packaging of the retroviral genome. Further biophysical characterization of RSV Gag-gRNA BMCs therefore possesses great potential to reveal novel avenues for therapeutic intervention.

## 1. Introduction

Effective control of retroviral infection requires a deeper understanding of the mechanisms underlying retroviral replication, due to their high mutation rate and associated potential for reduced efficacy of current therapies. As the primary structural component of retroviruses, Gag proteins represent an important target for study with high potential for novel discoveries of clinical significance.

The retroviral Gag protein directs virion assembly by selecting viral genomic RNA (gRNA) for incorporation into nascent virions [1–4]. Recognition of the viral genome is facilitated by a specific interaction between the nucleocapsid (NC) domain within Gag and the Ψ packaging signal near the 5’ end of the unspliced viral RNA (USvRNA) [5–7]. Mediated by two cis-acting elements within the USvRNA, the genome is selectively packaged into nascent virions as a noncovalently-linked dimer [3,8–14]. The first element is the dimerization initiation sequence (DIS), which forms a weak dimer via a kissing loop interaction. This dimer is then noncovalently stabilized by the dimerization linkage structure (DLS) [8,10,15–17]. Dimerization is absolutely required for infectivity and performs several functions, including facilitating strand transfer during reverse transcription and recombination between heterologous genome copies, leading to genetic diversity and repair of the sequence in the event of damage [13,18–21]. The association of Gag with gRNA triggers Gag multimerization, an important step in virion assembly and the production of infectious virus [22–24].

The dependence of retroviral Gag proteins on multimerization and nucleic acid binding for viral infectivity informed our recent line of investigation that identified the Gag proteins of both Rous sarcoma virus (RSV) and human immunodeficiency virus type-1 (HIV-1) as capable of forming biomolecular condensates (BMCs) *in vitro* and in cells [25–33]. It is increasingly appreciated that BMCs, phase-separated membraneless organelles driven by multivalent interactions between proteins and nucleic acids, are critical for many cellular functions [34–43]. In addition, numerous viruses have been shown to leverage viral and host BMCs for their replication [44–53]. In our studies, we observed the existence of Gag assemblies at multiple locations throughout the cell—in the nucleus, the cytoplasm, and at the plasma membrane—suggesting an important role for BMCs in retroviral particle assembly [26]. Notably, both RSV and HIV-1 Gag BMCs colocalized with their cognate USvRNAs at transcriptional burst sites in the nucleus, suggesting that co-transcriptional selection of gRNA by Gag may occur [1,2]. Our observation that RSV Gag preferentially packages genome homodimers further supports this hypothesis [13].

A key function of BMCs is to concentrate and compartmentalize protein and nucleic acid components essential for specific functions in the cell [34,35,38,39]. Based on our prior work, we hypothesized that retroviral Gag proteins form BMCs with genomic RNA to generate stable complexes that can remain intact as they travel through different subcellular compartments in transit to the plasma membrane for release of virions [1–3,13,26,27,33]. In support of this idea, our recent publication compared RSV Gag BMCs formed in the presence of vRNAs containing Ψ and/or the DIS to those formed with RNAs lacking these elements [54]. Notably, vRNAs containing Ψ had a stabilizing effect on RSV Gag BMCs which was further augmented compared to monomeric vRNAs by the addition of the DIS [54]. Furthermore, our data demonstrated that Gag BMCs could discriminate between Ψ and non-Ψ RNAs at physiologic salt concentrations, which was not observed in previous studies with Gag in solution, suggesting that using crowding agents to mimic the dense cellular environment may represent a more biologically relevant *in vitro* experimental system [54–56].

The most unexpected finding from our prior study was that a vRNA containing all of the elements necessary to form a stable dimer (Ψ, DIS, and DLS) resulted in the dramatic alteration of condensate architecture, producing Gag-vRNA BMCs that were bridged together by long strands of vRNA [54]. In contrast, monomeric Ψ-containing vRNAs resulted in discrete Gag-vRNA BMCs. Therefore, we sought to understand whether emergent biophysical properties imparted by this unique architecture could explain the mechanism(s) by which nanoscale Gag-gRNA complexes assemble into stable assemblies that facilitate gRNA packaging or macroscale assemblies that were observed by confocal microscopy. To this end, the present work utilized atomic force microscopy (AFM) and fluorescence correlation spectroscopy (FCS) approaches to determine the elasticity (AFM), diffusivity (FCS), and nanoscale morphologies (AFM) of Gag-vRNA BMCs [57–60]. Use of these techniques to study biological samples, including BMCs, has repeatedly demonstrated the power of such high resolution biophysical approaches in defining the spatial and mechanical properties that may contribute to biological function [57,58,61–71].

Our findings reveal a novel role for gRNA dimerization in nanoscale condensate architecture and stability that is consistent with the importance of gRNA dimerization for efficient packaging of the retroviral genome [14,18,19]. Further understanding how these characteristics of retroviral Gag-gRNA condensates contribute to efficient gRNA packaging and virus assembly has great potential to reveal novel avenues for therapeutic intervention.

## 2. Results

### 2.1. Biophysical Comparison of Similar Ψ-Containing Viral RNAs That Differ in Their Ability to Dimerize

Retroviral Gag proteins consist of multiple functional domains, each of which performs one or more role(s) in the assembly of virions (Figure 1A) [1–4]. A critical step in virus assembly is the selection and packaging of the dimeric retroviral genome, which is facilitated by a specific interaction between the NC domain and the Ψ packaging signal located near the 5’ end of USvRNA [3,5–14]. Recent work suggests that this selection event may occur co-transcriptionally, involving the entry of Gag into phase-separated transcriptional condensates [1,2,13,26,27,33,54]. The presence of positively-charged amino acids within the NC and matrix (MA) domains of Gag are critical for their roles in selecting gRNA and binding negatively-charged phospholipids in the plasma membrane, respectively, and also contribute to the nonspecific binding of cellular nucleic acids [3,72–78]. RSV and HIV-1 Gag proteins contain conformationally-promiscuous, intrinsically-disordered regions (IDRs) that may enable interaction with variable protein partners [26,27,54]. In RSV, IDRs exist within the MA, NC, p2, and p10 domains [26,54]. The ability of Gag proteins to bind a diverse set of proteins and nucleic acids promotes BMC formation.

**Figure 1.**
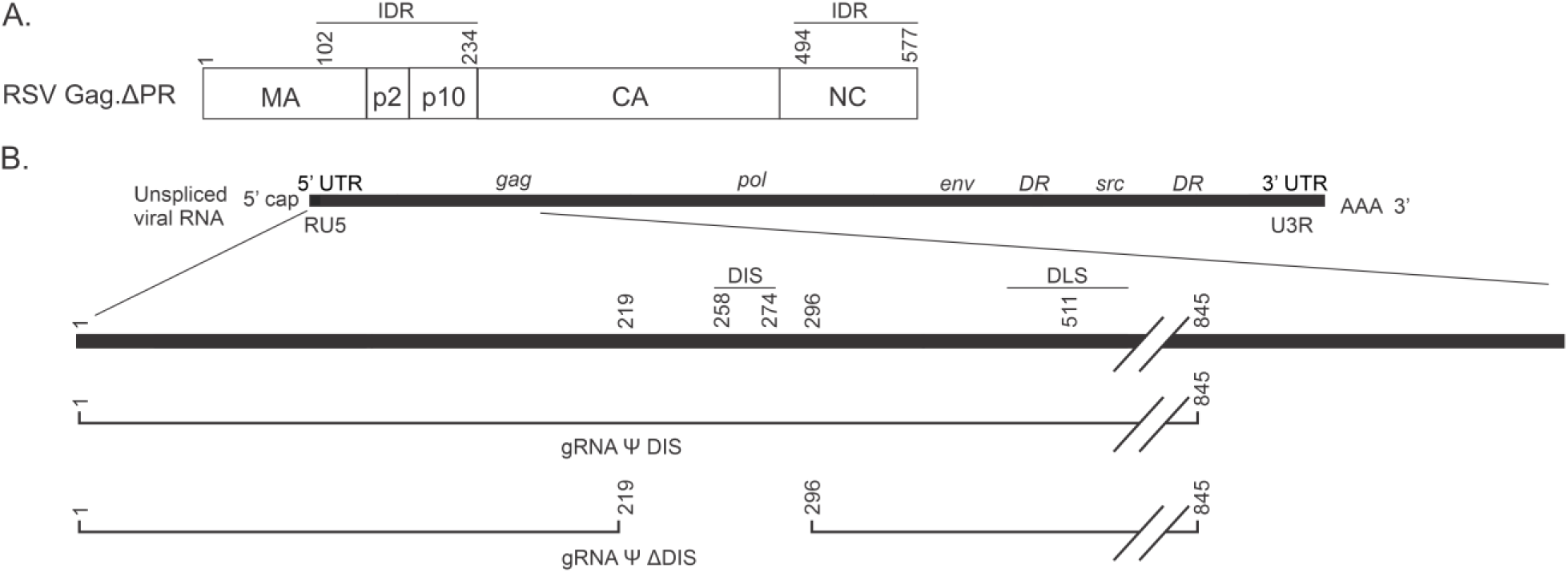
Schematics of RSV Gag.ΔPR and gRNA Constructs. (**A**) The recombinant RSV Gag.ΔPR protein contains the matrix (MA), p2, p10, capsid (CA), and nucleocapsid (NC) domains, but lacks the carboxy-terminal protease (PR) domain responsible for autoproteolysis of this multidomain protein. Recognition of the Ψ packaging sequence in the viral genomic RNA (gRNA) is mediated by the NC domain, and both the NC and MA domains nonspecifically interact with other nucleic acids. Two intrinsically-disordered regions (IDRs; amino acids ∼102-234 and ∼494-577) contribute to the formation of BMCs by this protein. (**B**) Schematic depicting the gRNA Ψ DIS and gRNA Ψ ΔDIS RNAs that are used in this manuscript. Regions encoding these RNAs are indicated in relation to the unspliced viral RNA from which they are derived; Ψ, psi packaging signal; DIS, dimerization initiation sequence; DLS, dimerization linkage structure. Figure adapted from Lambert et. al. 2025, PMID: 39861886.

We recently reported a comparison of the effect(s) of nonviral and viral RNAs on RSV Gag BMCs, revealing unique roles for viral RNAs containing Ψ and/or the DIS [54]. The most intriguing observation from that work was that a vRNA containing all of the elements necessary to form a stable dimer (Ψ, DIS, and DLS; Figure 1B, gRNA Ψ DIS) resulted in BMCs that were bridged together by long strands of vRNA [12,17,54,79,80]. This vRNA displayed high levels of colocalization with Gag and consistently promoted Gag BMC formation under a variety of assay conditions. Notably, a similar vRNA containing Ψ but lacking the DIS (Figure 1B, gRNA Ψ ΔDIS) and therefore the ability to dimerize, resulted in discrete Gag-vRNA BMCs, lower levels of colocalization, and much more variable effects on condensate abundance [54]. These findings complement our previous work demonstrating preferential packaging of RSV gRNA homodimers and support a role for BMCs in the preferential selection of gRNA dimers [13,54]. We therefore pursued the question of how dynamic structures such as BMCs remain mechanically and compositionally stable during transit from the nucleus to the plasma membrane.

The use of high-resolution biophysical approaches such as AFM and FCS to define the architectural and mechanical properties that may contribute to biological function is well documented [57,58,61–71]. Therefore, we sought to utilize these approaches to understand if emergent biophysical properties imparted by this unique architecture could explain the mechanism(s) by which nanoscale Gag-gRNA complexes form stable BMCs that provide selectivity and facilitate gRNA packaging. To achieve this goal, we compared the two similar vRNAs described above (Figure 1B) in regards to their effect(s) on condensate architecture and resistance to deformation at the nanoscale using AFM. We then probed the diffusion of Gag and RNA molecules within these BMCs using FCS.

### 2.2. Assessing the Nanoscale Architecture of RSV Gag-vRNA Co-Condensates

As mentioned above, striking differences in condensate architecture were observed between dimerization-competent gRNA Ψ DIS and monomeric gRNA Ψ ΔDIS via confocal fluorescence imaging in our prior studies [54]. Despite its many strengths and our utilization of multiple advanced image processing and analysis techniques available to us, confocal imaging still has resolution limits that may preclude our ability to discern fine morphological differences between Gag-vRNA BMCs. As such, we utilized AFM to characterize the morphology of BMCs formed either by RSV Gag alone, or in complex with either of the two vRNAs described above (Figure 2). To aid in the comparison of BMC morphologies between confocal and AFM approaches, we are showing new examples of condensates at both lower and higher magnification from our previously-acquired confocal images to demonstrate the unique morphology of Gag-gRNA Ψ DIS BMCs versus Gag-only and Gag-gRNA Ψ ΔDIS BMCs (Figure 2A, C, E). AFM imaging further refined these distinctions, revealing subtle differences between Gag alone and Gag-gRNA Ψ ΔDIS BMCs, in addition to confirming the linked structure of Gag BMCs formed with dimeric gRNA Ψ DIS (Figure 2B, D, F). The somewhat smoother and less ridged morphology of Gag-gRNA Ψ ΔDIS BMCs compared to Gag alone BMCs is likely reflective of the contribution of RNA to the overall condensate structure (Figure 2B, F). Intriguingly, AFM also revealed that the “peapod-like” vRNA shell observed with gRNA Ψ DIS via confocal microscopy more tightly surrounds the regions containing Gag than that approach would suggest (Figure 2C-D). The fundamental morphological differences observed between Gag-vRNA condensates in the presence or absence of dimerization-competent vRNA potentially suggests a biomechanical role in these condensates.

**Figure 2.**
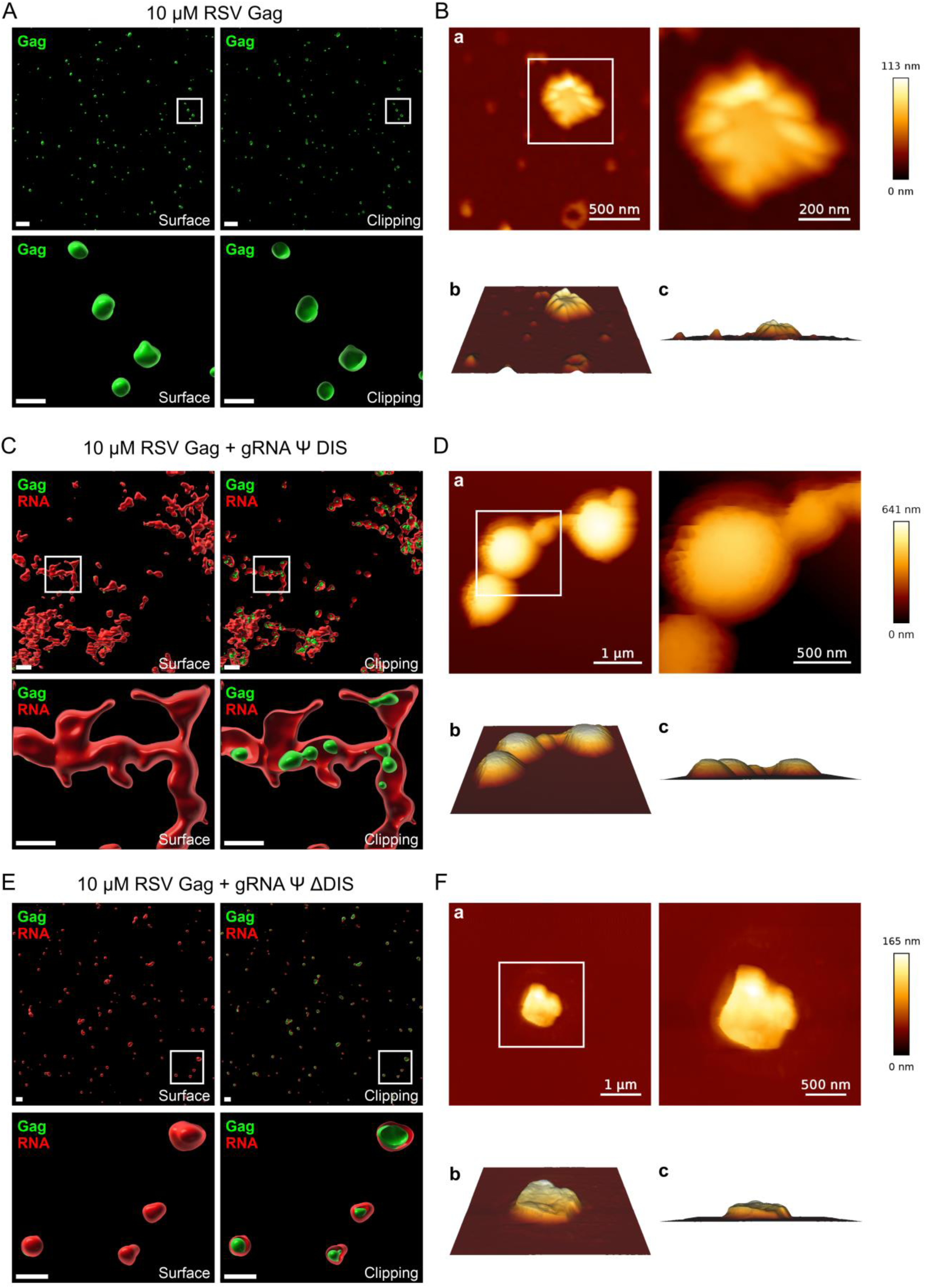
Confocal and atomic force microscopy analysis of RSV Gag-vRNA co-condensate morphology. Condensate morphology was assessed using both confocal (**A**, **C**, **E**) and atomic force microscopy (**B**, **D**, **F**) for RSV Gag alone (**A**-**B**), RSV Gag + gRNA Ψ DIS (**C**-**D**), and RSV Gag + gRNA Ψ ΔDIS (**E**-**F**). For fluorescence microscopy images (**A**, **C**, **E**), raw images were deconvolved and used to generate three-dimensional surfaces in Imaris Image Analysis Software. Both surface renderings (left panels) and orthogonal clipping plane images (right panels) are depicted. Regions boxed in the top set of images are enlarged below. Scale bars are either 2 μm (whole field, top panels) or 1 μm (enlarged boxed regions, bottom panels). For atomic force microscopy images (**B**, **D**, **F**), regions boxed in (**a**) are enlarged to the right. Three-dimensional renderings at 60° and 90° (**b** and **c**, respectively) are depicted below. Scale bars are as indicated.

### 2.3. Viral RNA Dimerization Promotes the Formation of Heterogeneous Condensate Assemblies Displaying Increased Resistance to Mechanical Deformation

Having verified a role for gRNA Ψ DIS dimers in a unique nanoscale Gag-vRNA co-condensate architecture compared to Gag alone and Gag-gRNA Ψ ΔDIS, we next wanted to leverage AFM to probe the biomechanical properties of these condensates. More specifically, we wanted to assess if the contribution of RNA dimers within these BMCs might confer mechanical stability that could help explain the preferential packaging of gRNA dimers in light of potential intermolecular and mechanical interference during trafficking from the nucleus to the plasma membrane. This was achieved by utilizing an AFM force spectroscopy modality to simultaneously image condensate morphology and assess the resistance of said condensates to deformation by determination of the elastic modulus, also known as Young’s Modulus (Figure 3). This approach revealed a number of interesting observations, the most striking of which was that dimeric gRNA Ψ DIS promotes the formation of BMCs with heterogeneous mechanical properties, including nanoclusters with greatly increased resistance to mechanical deformation (Figure 3A, C). Notably, these nanoclusters appear to be linked by regions of intermediate resistance to deformation, and both of these regions appear to be further surrounded by a much more deformable area. This is consistent with the idea of Gag-gRNA Ψ DIS nanoclusters joining together via inter-condensate dimerization, producing regions of intermediate elasticity prior to coalescence. The even less elastic regions would then represent areas dominated by strands of vRNA available for interaction with disparate BMCs. The multimodal distribution of Young’s Modulus values observed for the BMCs in Figure 3A is reflective of the existence of these different regions. In contrast, BMCs formed under the remaining two conditions exhibited more homogenous mechanical properties with a bias towards more deformable structures. This property posed challenges in the accurate determination of Young’s Modulus at higher resolutions, and is therefore the reason for the variability in the number of pixels sampled in Figure 3B-D.

**Figure 3.**
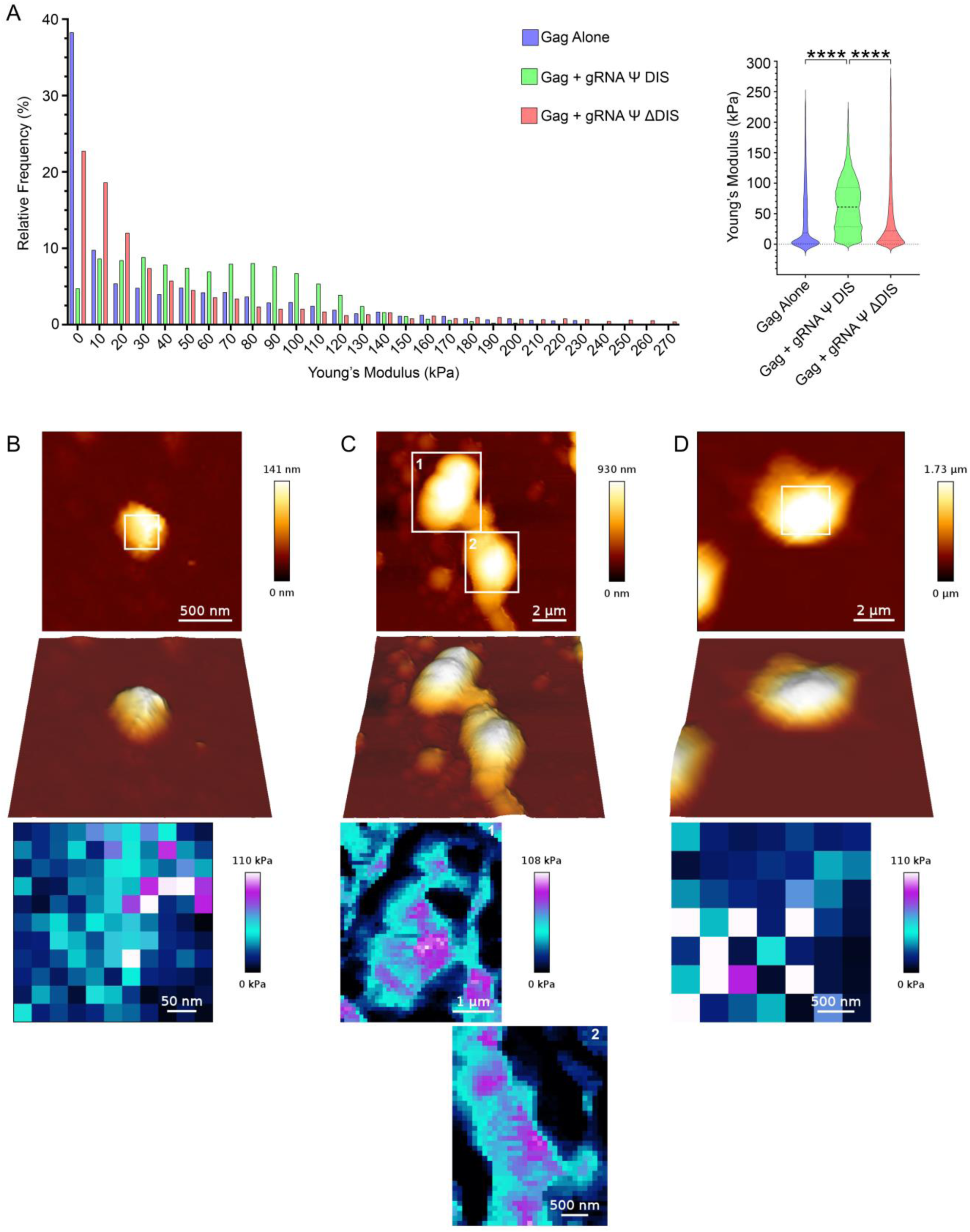
RSV Gag-gRNA Ψ DIS co-condensates contain nanoclusters with increased resistance to mechanical deformation. (**A**) Elasticity (Young’s Modulus, kPa) values for Gag and Gag-vRNA condensates are depicted in histogram (left; bin size = 10, bin center plotted) and violin plot (right; median, first, and third quartile values displayed by dotted lines) form. Statistical significance was determined by Kruskal–Wallis test with Dunn’s post-hoc test (****, p ≤ 0.0001), n ≥ 3849. (**B**-**D**) Representative atomic force microscopy images (top) and modulus maps corresponding to boxed regions (bottom). Scale bars are as indicated.

### 2.4. Viral RNA Dimerization Results in the Formation of RNA Scaffolds

To determine whether the marked difference in mechanical stability observed in the presence of dimeric gRNA Ψ DIS was accompanied by a difference in diffusivity of Gag and/or RNA molecules, we utilized FCS to measure the diffusion coefficients of each of these species at multiple locations within each BMC (Center, C; Intermediate, I; Edge, E; Figure 4 and Table 1). At physiologic salt concentrations (150 mM), we observed that BMCs formed by Gag alone were dynamically arrested at all locations within BMCs, as evidenced by extremely low mean diffusion coefficients (C: 0.423 µm^2^/s; I: 0.429 µm^2^/s; E: 0.363 µm^2^/s)(Figure 4A). This observation was consistent with our previous observation that Gag self-assembles into stable complexes at this salt concentration that demonstrate limited mobility by fluorescence recovery after photobleaching (FRAP) [54]. However, the addition of either monomeric or dimeric vRNAs significantly increased mean Gag diffusivity in each region of the condensate [gRNA Ψ DIS (C: 48.3 µm^2^/s; I: 50.5 µm^2^/s; E: 87.2 µm^2^/s) versus gRNA Ψ ΔDIS (C: 35.4 µm^2^/s; I: 40.2 µm^2^/s; E: 86.3 µm^2^/s)](Figure 4A). Strikingly, both vRNAs displayed similarly low diffusivity independent of location within BMCs [gRNA Ψ DIS (C: 0.209 µm^2^/s; I: 0.268 µm^2^/s; E: 0.299 µm^2^/s) compared to gRNA Ψ ΔDIS (C: 0.205 µm^2^/s; I: 0.247 µm^2^/s; E: 0.244 µm^2^/s)](Figure 4B). Representative images are displayed in Figure 4C, again demonstrating the morphological differences between BMCs formed in the presence of these two vRNAs at 150 mM NaCl.

**Figure 4.**
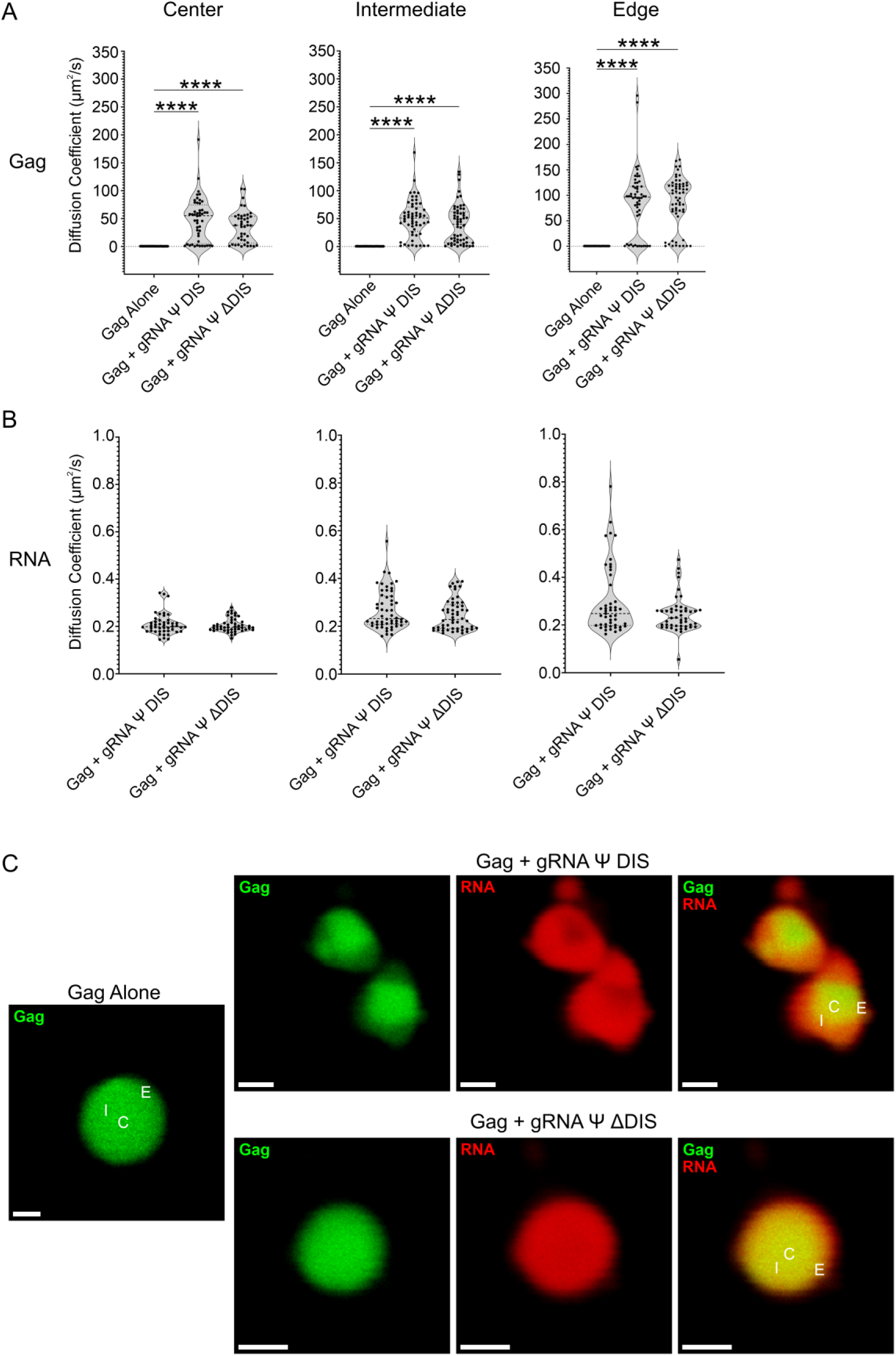
Fluorescence correlation spectroscopy (FCS) diffusivity analysis of RSV Gag-vRNA condensates at 150 mM NaCl. (**A**, **B**) Violin plots containing Gag (**A**) and RNA (**B**) diffusion coefficient (μm^2^/s) values for Gag alone, Gag-gRNA Ψ DIS, and Gag-gRNA Ψ ΔDIS condensates as determined by FCS. Diffusivity was analyzed at three independent locations within condensates. Median, first, and third quartile values are displayed by dotted lines. Statistical significance was determined by Kruskal–Wallis test with Dunn’s post-hoc test (**A**) (****, p ≤ 0.0001) or Mann–Whitney test (**B**) (n ≥ 42 for all datasets). (**C**) Representative images of Gag alone, Gag-gRNA Ψ DIS, and Gag-gRNA Ψ ΔDIS condensates analyzed by FCS. Scale bars = 1 μm. C, center; I, intermediate; E, edge.

**Table 1.** Diffusivity of RSV Gag Alone or Gag-vRNA Condensates.

| Condition | Gag Diffusion Coefficient ($\mu\text{m}^2/\text{s}$) | RNA Diffusion Coefficient ($\mu\text{m}^2/\text{s}$) |
| --- | --- | --- |
| Gag Alone: Center, 150 mM | 0.423 | N/A |
| Gag Alone: Intermediate, 150 mM | 0.429 | N/A |
| Gag Alone: Edge, 150 mM | 0.363 | N/A |
| gRNA $\Psi$ DIS: Center, 150 mM | 48.3 | 0.209 |
| gRNA $\Psi$ DIS: Intermediate, 150 mM | 50.5 | 0.268 |
| gRNA $\Psi$ DIS: Edge, 150 mM | 87.2 | 0.299 |
| gRNA $\Psi$ ΔDIS: Center, 150 mM | 35.4 | 0.205 |
| gRNA $\Psi$ ΔDIS: Intermediate, 150 mM | 40.2 | 0.247 |
| gRNA $\Psi$ ΔDIS: Edge, 150 mM | 86.3 | 0.244 |
| Gag Alone: Center, 500 mM | 40.4 | N/A |
| Gag Alone: Intermediate, 500 mM | 47.1 | N/A |
| Gag Alone: Edge, 500 mM | 78.5 | N/A |
| gRNA $\Psi$ DIS: Center 500 mM | 17.1 | 0.217 |
| gRNA $\Psi$ DIS: Intermediate, 500 mM | 32.7 | 0.226 |
| gRNA $\Psi$ DIS: Edge, 500 M | 89.0 | 0.252 |
| gRNA $\Psi$ ΔDIS: Center 500 mM | 30.8 | 0.228 |
| gRNA $\Psi$ ΔDIS: Intermediate, 500 mM | 43.9 | 0.239 |
| gRNA $\Psi$ ΔDIS: Edge, 500 mM | 78.5 | 0.265 |
| Gag Alone: Center, 1 M | 39.9 | N/A |
| Gag Alone: Intermediate, 1 M | 27.1 | N/A |
| Gag Alone: Edge, 1 M | 68.3 | N/A |
| gRNA $\Psi$ DIS: Center 1 M | 30.6 | 0.260 |
| gRNA $\Psi$ DIS: Intermediate, 1 M | 32.0 | 0.249 |
| gRNA $\Psi$ DIS: Edge, 1 M | 71.7 | 0.215 |
| gRNA $\Psi$ ΔDIS: Center 1 M | 16.8 | 0.249 |
| gRNA $\Psi$ ΔDIS: Intermediate, 1 M | 20.7 | 0.266 |
| gRNA $\Psi$ ΔDIS: Edge, 1 M | 73.1 | 0.244 |

We previously demonstrated that the RSV Gag-Ψ interaction is resistant to ionic disruption within BMCs [54]. Therefore, we also probed Gag and vRNA diffusivity in BMCs at higher salt concentrations (500 mM and 1 M NaCl) to assess the contribution of electrostatics to this property (Figures 5, 6 and Table 1). At 500 mM NaCl, we observed an increase in mean diffusivity in Gag-only BMCs, signaling a weakening of Gag-Gag interactions at this salt concentration (C: 40.4 µm^2^/s; I: 47.1 µm^2^/s; E: 78.5 µm^2^/s)(Figure 5A). Under these conditions, neither vRNA further increased Gag diffusivity [gRNA Ψ DIS (C: 17.1 µm^2^/s; I: 32.7 µm^2^/s; E: 89.0 µm^2^/s), gRNA Ψ ΔDIS (C: 30.8 µm^2^/s; I: 43.9 µm^2^/s; E: 78.5 µm^2^/s)](Figure 5A). In fact, Gag diffusivity at the center of Gag-gRNA Ψ DIS BMCs was significantly decreased compared to Gag or Gag-gRNA Ψ ΔDIS BMCs.

**Figure 5.**
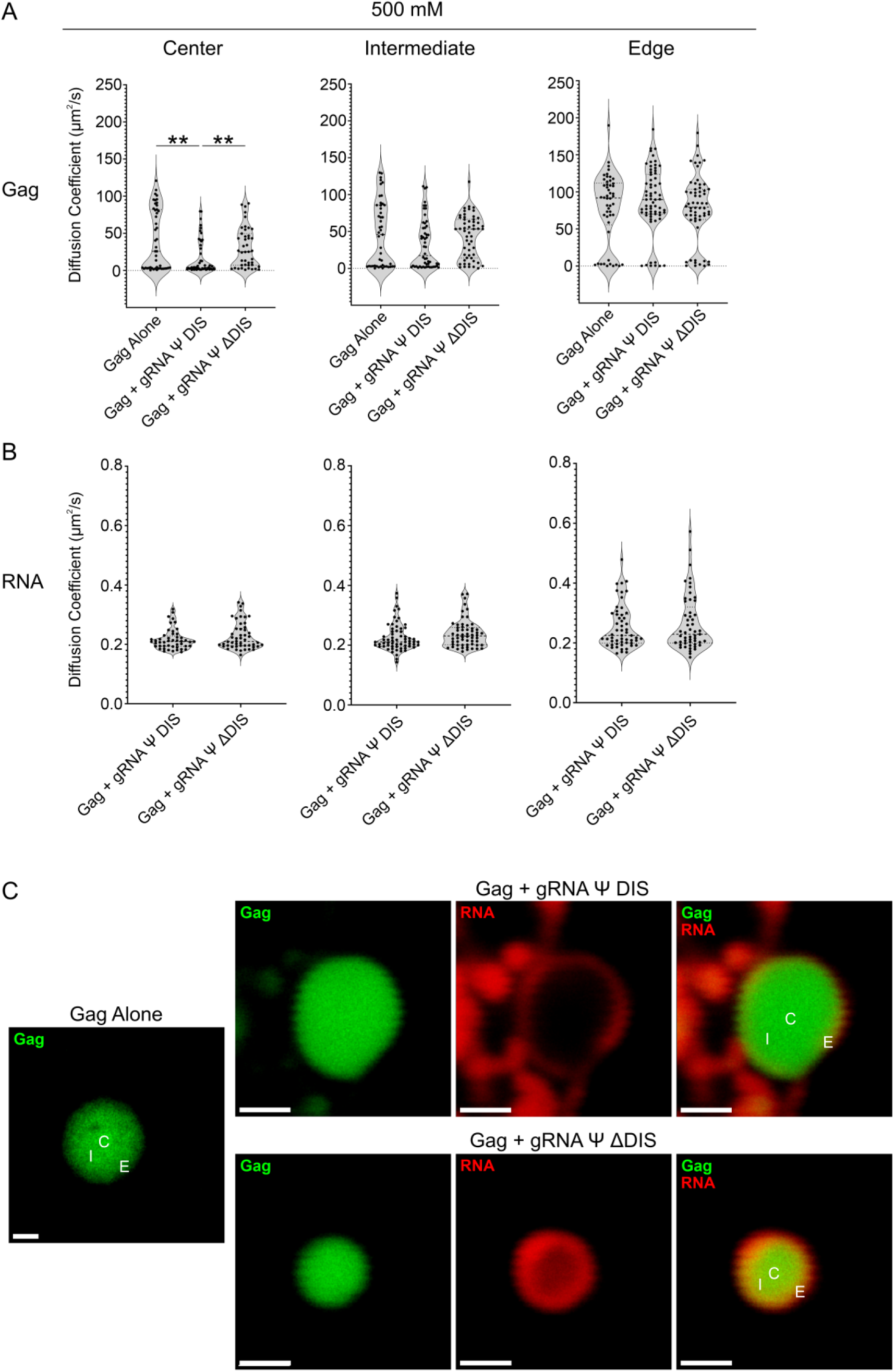
Fluorescence correlation spectroscopy (FCS) diffusivity analysis of RSV Gag-vRNA condensates at 500 mM NaCl. (**A**, **B**) Violin plots containing Gag (**A**) and RNA (**B**) diffusion coefficient (μm^2^/s) values for Gag alone, Gag-gRNA Ψ DIS, and Gag-gRNA Ψ ΔDIS condensates as determined by FCS. Diffusivity was analyzed at three independent locations within condensates. Median, first, and third quartile values are displayed by dotted lines. Statistical significance was determined by Kruskal–Wallis test with Dunn’s post-hoc test (**A**) (**, p ≤ 0.01) or Mann–Whitney test (**B**) (n ≥ 43 for all datasets). (**C**) Representative images of Gag alone, Gag-gRNA Ψ DIS, and Gag-gRNA Ψ ΔDIS condensates analyzed by FCS. Scale bars = 1 μm. C, center; I, intermediate; E, edge.

**Figure 6.**
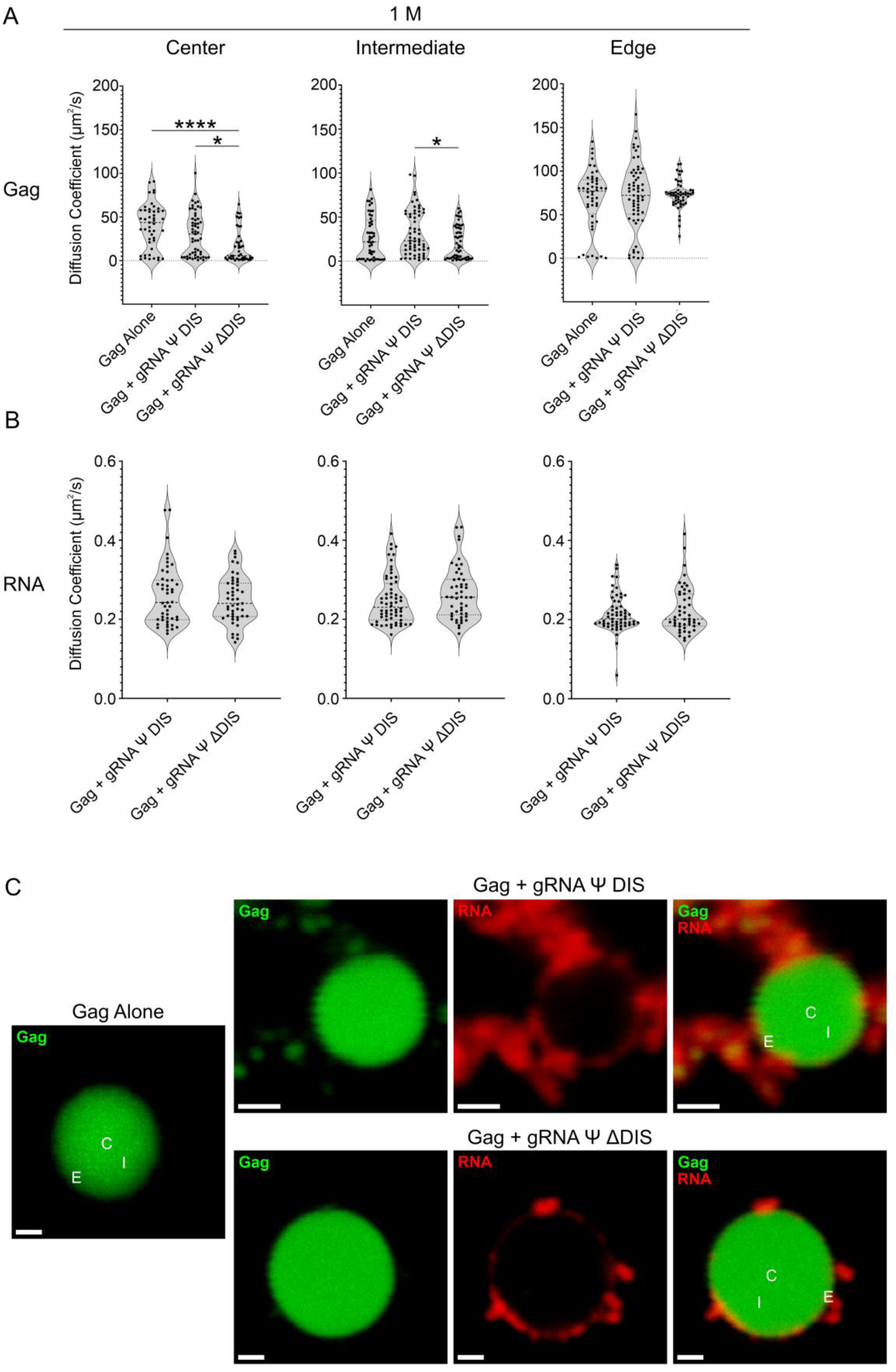
Fluorescence correlation spectroscopy (FCS) diffusivity analysis of RSV Gag-vRNA condensates at 1 M NaCl. (**A**, **B**) Violin plots containing Gag (**A**) and RNA (**B**) diffusion coefficient (μm^2^/s) values for Gag alone, Gag-gRNA Ψ DIS, and Gag-gRNA Ψ ΔDIS condensates as determined by FCS. Diffusivity was analyzed at three independent locations within condensates. Median, first, and third quartile values are displayed by dotted lines. Statistical significance was determined by Kruskal–Wallis test with Dunn’s post-hoc test (**A**) (****, p ≤ 0.0001; *, p ≤ 0.05) or Mann–Whitney test (**B**) (n ≥ 41 for all datasets). (**C**) Representative images of Gag alone, Gag-gRNA Ψ DIS, and Gag-gRNA Ψ ΔDIS condensates analyzed by FCS. Scale bars = 1 μm. C, center; I, intermediate; E, edge.

Values for vRNA diffusivity were not affected by 500 mM salt, remaining extremely low [gRNA Ψ DIS (C: 0.217 µm^2^/s; I: 0.226 µm^2^/s; E: 0.252 µm^2^/s), gRNA Ψ ΔDIS (C: 0.228 µm^2^/s; I: 0.239 µm^2^/s; E: 0.265 µm^2^/s)(Figure 5B). Representative images shown in Figure 5C demonstrate a partial exclusion of both RNAs from the center of Gag-vRNA BMCs, although this displacement was more pronounced for gRNA Ψ DIS.

At 1 M NaCl, there was again no significant increase in diffusivity in the presence of either RNA compared to Gag alone [Gag alone (C: 39.9 µm^2^/s; I: 27.1 µm^2^/s; E: 68.3 µm^2^/s), gRNA Ψ DIS (C: 30.6 µm^2^/s; I: 32.0 µm^2^/s; E: 71.7 µm^2^/s), gRNA Ψ ΔDIS (C: 16.8 µm^2^/s; I: 20.7 µm^2^/s; E: 73.1 µm^2^/s)], nor was there any substantial change in the diffusivity of these two RNAs within BMCs [gRNA Ψ DIS (C: 0.260 µm^2^/s; I: 0.249 µm^2^/s; E: 0.215 µm^2^/s), gRNA Ψ ΔDIS (C: 0.249 µm^2^/s; I: 0.266 µm^2^/s; E: 0.224 µm^2^/s)(Figure 6A-B). Interestingly, there was no longer a decrease in Gag diffusivity at the center of Gag-gRNA Ψ DIS BMCs, with Gag-gRNA Ψ ΔDIS BMCs instead now displaying decreased diffusivity at this location. Representative images of these two types of BMCs revealed that both RNAs were largely excluded from the condensate center (Figure 6C).

## 3. Discussion

A better understanding of the mechanisms underlying retroviral replication has great potential to identify novel targets for antiviral therapy. To this end, our work has resulted in important revelations that have improved the understanding of the mechanisms underlying Gag-gRNA packaging and retrovirus assembly. The observations that Gag proteins undergo transient nucleocytoplasmic trafficking [81–83], colocalize with USvRNA at sites of active transcription [1,2], and form BMCs via IDR- and electrostatic-mediated interactions (in addition to the specific interaction with Ψ) [26,27,33,54], provide a basis for answering key questions in the field while simultaneously inspiring new lines of investigation. At the heart of this work are questions concerning the role of BMCs in retroviral gRNA selection and the successful trafficking of Gag-gRNA complexes to the plasma membrane for virion assembly.

The present work stems from previous observations that gRNA Ψ DIS, a viral RNA containing all of the elements needed to form a stable dimer (Ψ, DIS, and DLS), forms heterogeneous condensates that display a “peapod-like” morphology via confocal microscopy wherein Gag-rich BMCs (“peas”) are contained within and linked together via an RNA “pod” formed by inter-condensate RNA dimers [54]. This architecture was unique among all of the RNAs we tested in our prior study, leading us to postulate that gRNA dimerization may serve another role in the promotion of viral fitness in addition to those related to genetic diversity and genome fidelity [13,18–21]. One intriguing possibility is that gRNA dimerization works in concert with the Gag-Ψ interaction to confer selectivity within transcriptional condensates. In addition to the well-known nucleic acid binding properties of NC and MA, proteomic studies suggest that Gag may interact with a variety of host transcription-related proteins [33,55,56,72–74,84–89]. These interactions are likely important for participation in transcriptional BMCs, but they must eventually be overcome for the Gag-gRNA complex to be released from the transcription site. The mechanism mediating compositional selectivity may be dependent on dimer-mediated nucleation of discrete Gag-gRNA complexes into larger assemblies possessing emergent biophysical properties. Precedence for this concept exists in *Drosophila melanogaster*, where the kissing-loop homodimerization of *bicoid* and *oskar* RNAs is important for the nucleation of compositionally-distinct BMCs with the RNA binding proteins Staufen and Bruno [90–93]. The notion that Gag-gRNA BMCs may be functionally similar to those from Drosophila is intriguing, given that their major function is the packaging and transport of these RNAs [90]. Further support for the above possibility is evident in the unique architecture and biomechanical heterogeneity of Gag-gRNA Ψ DIS BMCs (Figures 2 and 3), and the observation that this vRNA both robustly promoted condensate formation and displayed high levels of colocalization in our prior study [54].

Another intriguing result of the work contained herein is that Gag-gRNA Ψ DIS BMCs display emergent biophysical properties, as determined by AFM and FCS approaches (Figures 3, 4, 5, and 6). As has been demonstrated with cellular structures, utilization of AFM to probe the biomechanical properties of biological samples can reveal features that are not otherwise discernible [68]. Analyzing Gag-vRNA BMCs in this manner revealed underlying regions of increased elasticity that likely correspond to stable Gag-vRNA nanoclusters (Figure 3A, C). Notably, these nanoclusters appear to be linked by regions of intermediate resistance to deformation, likely composed of less stable Gag-vRNA complexes, which are then further surrounded by a much more deformable area that is likely RNA-rich. This arrangement is again in good agreement with the evidence for gRNA dimer-driven nucleation discussed above.

This heterogeneity in biophysical properties was likewise evident by FCS (Figures 4, 5, 6), with the multimodal nature of Gag diffusivity values reflective of the multiple populations of Gag described above. Interestingly, the relative immobility of RNA molecules in these assays parallels that of Whi3-RNA condensates from *Ashbya gossypii*, where dynamical arrest of RNA within the condensates promotes their retention and maintains condensate composition in the presence of RNAs that also form BMCs with Whi3 [94,95]. Dynamical arrest is dependent on RNA scaffolding driven by secondary structure and sequence complementarity [95]. Such scaffolding was also shown to be resistant to disruption by high salt conditions, which we also observed in our FCS experiments (Figures 5 and 6) [94]. Therefore, the likelihood that a similar mechanism is at work in Gag-gRNA condensates is high.

This study utilized advanced imaging and biophysical techniques to characterize RSV Gag-vRNA co-condensates, revealing that the presence of dimeric vRNA results in both a unique nanoscale condensate architecture and biophysical profile. Taken together, these data suggest that the formation of compositionally- and biomechanically-stable Gag-gRNA condensates is likely a critical step in the assembly of infectious virions, facilitating the efficient selection, trafficking, and packaging of the dimeric retroviral genome. As has been proposed for other condensate-utilizing viral systems, drug-mediated disruption of Gag-gRNA BMCs represents a novel approach for the prevention of virion assembly [49,50,96]. Therefore, further characterization of the fundamental processes driving the formation and function of retroviral Gag-gRNA condensates has great potential to identify mechanisms by which to disrupt these interactions for clinical benefit.

## 4. Materials and Methods

### 4.1. Plasmids

The plasmid encoding the RSV Gag protein used for purification [pET28(-His) Gag.ΔPR (henceforth referred to as RSV Gag)] was described in [97]. Plasmids for production of viral RNA were generated by cloning the sequences encoding gRNA Ψ DIS (also known as 1–845 Ψ DIS [17,54,79]) and gRNA Ψ ΔDIS (also known as 1–219/296–845 Ψ ΔDIS [17,54,79]) into a pGEM vector backbone, which were then used to generate PCR templates encoding gRNA Ψ DIS and gRNA Ψ ΔDIS using the same forward (5′ ATCGTAATACGACTCACTATAGGCCATTTGACCATTCACCACATTGGTG) and reverse (TATCGATTTCGAACCCGGGGTACC) primers. These PCR templates were then used to generate the desired RNA via *in vitro* transcription reactions.

### 4.2. Gag Protein Expression, Purification, and Fluorescent Labeling

Recombinant RSV Gag was expressed and purified, as described previously [26,98], and was labeled using an Alexa Fluor 488 Microscale Protein Labeling Kit (Thermo Fisher Scientific, Waltham, MA, USA, #A30006) according to the manufacturer’s instructions. Protein concentration was quantified via NanoDrop One spectrophotometer (Thermo Fisher Scientific, Waltham, MA, USA), and was then aliquoted into working volumes, flash-frozen in dry ice/ethanol, and stored at −80 °C until use.

### 4.3. RNA Synthesis by In Vitro Transcription and Fluorescent Labeling

The PCR products mentioned above were used as template to generate RNAs using a MEGAscript T7 Transcription Kit (Thermo Fisher Scientific, Waltham, MA, USA, #AM1334) according to the manufacturer’s instructions. In order to fluorescently label RNAs, aminoallyl-UTP was included in the reaction at a ratio of 4:1 to unmodified UTP. RNA was subsequently cleaned up via an RNA Clean and Concentrator Kit (Zymo Research Corp., Irvine, CA, USA, #R1014), and length and purity were assessed via agarose gel electrophoresis. RNAs were then labeled using an Alexa Fluor 647 Reactive Dye Decapack (Invitrogen, Waltham, MA, USA, #A32757) per the manufacturer’s instructions, and concentration was determined via a NanoDrop One spectrophotometer (Thermo Fisher Scientific, Waltham, MA, USA). RNA was aliquoted into working volumes, flash-frozen in dry ice/ethanol, and stored at −80 °C until use.

### 4.4. RNA Sequences

gRNA Ψ DIS: 5′ GCCAUUUGACCAUUCACCACAUUGGUGUGCACCUGGGUUGAUGGCCGGACCGUUGAUUCCCUGACGACUACGAGCACCUGCAUGAAGCAGAAGGCUUCAUUUGGUGACCCCGACGUGAUAGUUAGGGAAUAGUGGUCGGCCACAGACGGCGUGGCGAUCCUGUCUCCAUCCGUC UCGUCUAUCGGGAGGCGAGUUCGAUGACCCUGGUGGAGGGGGCUGCGGCUUAGGGAGGCAGAAGCUGAGUACCGUCGGAGGGAGCUCUACUGCAGGGAGCCCAGAUACCCUACCGAGAACUCAGAGAGUCGUUGGAAGACGGGAAGGAAGCCCGACGACUGAGCAGUCCACCCCAGGCGUGAUUCUGGUCGCCCGGUGGAUCAAGCAUGGAAGCCGUCAUAAAGGUGAUUUCGUCCGCGUGUAAAACCUAUUGCGGGAAAACCUCUCCUUCUAAGAAGGAAAUAGGGGCCAUGUUGUCCCUCUUACAAAAGGAAGGGUUGCUUAUGUCUCCCUCAGACUUAUAUUCCCCGGGGUCCUGGGAUCCCAUUACCGCGGCGCUAUCCCAGCGGGCUAUGAUACUUGGGAAAUCGGGAGAGUUAAAAACCUGGGGAUUGGUUUUGGGGGCAUUGAAGGCGGCUCGAGAGGAACAGGUUACAUCUGAGCAAGCAAAGUUUUGGUUGGGAUUAGGGGGAGGGAGGGUCUCUCCCCCAGGUCCGGAGUGCAUCGAGAAACCAGCAACGGAGCGGCGAAUCGACAAAGGGGAGGAAGUGGGAGAAACAACUGUGCAGCGAGAUGCGAAGAUGGCGCCGGAGGAAACGGCCACACCUAAAACCGUUGGCACAUCCUGCUAU 3′ gRNA Ψ ΔDIS: 5′ GCCAUUUGACCAUUCACCACAUUGGUGUGCACCUGGGUUGAUGGCCGGACCGUUGAUUCCCUGACGACUACGAGCACCUGCAUGAAGCAGAAGGCUUCAUUUGGUGACCCCGACGUGAUAGUUAGGGAAUAGUGGUCGGCCACAGACGGCGUGGCGAUCCUGUCUCCAUCCGUCUCGUCUAUCGGGAGGCGAGUUCGAUGACCCUGGUGGAGGGGGCUAGAGAGUCGUUGGAAGACGGGAAGGAAGCCCGACGACUGAGCAGUCCACCCCAGGCGUGAUUCUGGUCGCCCGGUGGAUCAAGCAUGGAAGCCGUCAUAAAGGUGAUUUCGUCCGCGUGUAAAACCUAUUGCGGGAAAACCUCUCCUUCUAAGAAGGAAAUAGGGGCCAUGUUGUCCCUCUUACAAAAGGAAGGGUUGCUUAUGUCUCCCUCAGACUUAUAUUCCCCGGGGUCCUGGGAUCCCAUUACCGCGGCGCUAUCCCAGCGGGCUAUGAUACUUGGGAAAUCGGGAGAGUUAAAAACCUGGGGAUUGGUUUUGGGGGCAUUGAAGGCGGCUCGAGAGGAACAGGUUACAUCUGAGCAAGCAAAGUUUUGGUUGGGAUUAGGGGGAGGGAGGGUCUCUCCCCCAGGUCCGGAGUGCAUCGAGAAACCAGCAACGGAGCGGCGAAUCGACAAAGGGGAGGAAGUGGGAGAAACAACUGUGCAGCGAGAUGCGAAGAUGGCGCCGGAGGAAACGGCCACACCUAAAACCGUUGGCACAUCCUGCUAU 3′

### 4.5. In Vitro Condensate Formation

Condensates were formed *in vitro*, as described previously [54]. Briefly, the following buffers were utilized: (i) No Salt Buffer: 0 mM NaCl, 50 mM HEPES pH 7.5, 1 mM DTT, 1 mM PMSF; (ii) High Salt Buffer: 4000 mM NaCl, 50 mM HEPES pH 7.5, 1 mM DTT, 1 mM PMSF; (iii) Crowding Buffer: 50% w/v polyethylene glycol-8000 (PEG-8000) in No Salt Buffer. Prior to use, the Crowding Buffer was warmed in a 42 °C water bath with periodic mixing for at least 1 h to ensure PEG-8000 was homogenously dissolved. No Salt and High Salt buffers were combined in Low-Binding microcentrifuge tubes (VWR, West Chester, PA, USA, #76332-066) to achieve the desired final NaCl concentration once all components had been added (protein, RNA, and Crowding Buffer).

RSV Gag proteins (unlabeled and Alexa Fluor 488-labeled) were thawed to room temperature, centrifuged at 21,000 × *g* for 2 min to pellet insoluble material, and transferred to fresh tubes. RSV Gag protein was added to the reaction tubes at a ratio of 1 (labeled) to 10 (unlabeled). RNA was then added, and the reaction was mixed via gentle pipetting. Crowding Buffer was added to a final concentration of 10% w/v and the reaction was thoroughly mixed. Reactions were incubated at room temperature for 1 h before use.

### 4.6. Confocal Imaging and Three-Dimensional Reconstructions

For confocal imaging, 6 μL of the reaction was deposited on a glass coverslip and a glass slide was placed on top. After 5 min, the coverslip was sealed with nail polish. Slides were then imaged using a Leica AOBS SP8 FALCON confocal microscope with a 63x/1.2 water objective (Leica Microsystems, Wetzlar, Germany). White light laser (WLL) line excitations of 488 and 647 nm were used in conjunction with hybrid detectors to detect 488-labeled protein and 647-labeled RNA, respectively. A minimum of three z-stacks were acquired at 3X and 7X zoom per condition for three-dimensional analyses.

To make three-dimensional reconstructions, deconvolved images were analyzed in Imaris Image Analysis Software v10.2.0 (Oxford Instruments, Abingdon, United Kingdom) using 3X and 7X zoom z-stacks. Surface renderings were generated from the Gag and RNA signals, and a surface image was taken. The orthogonal clipping plane tool was used to view the internal architecture of the condensates, and a clipping image was taken to demonstrate the spatial arrangement of Gag and RNA in condensates.

### 4.7. Atomic Force Microscopy (AFM)

For AFM imaging, 30 μL of condensate reactions were deposited on freshly cleaved mica surfaces mounted to glass slides. After 30 minutes, mica surfaces were rinsed three times with reaction buffer (150 mM NaCl, 50 mM HEPES pH 7.5, 1 mM DTT, 1 mM PMSF) to remove unbound condensates before imaging. A Bruker NanoWizard® Ultra Speed 2 atomic force microscope (Bruker, Billerica, MA, USA) was utilized to image condensates in either PeakForce Tapping (morphology) or Quantitative Imaging (Young’s Modulus) mode. Bruker BioAFM Data Processing software v8.0 (Bruker, Billerica, MA, USA) was used to analyze and export images and to quantify Young’s Modulus data. Modulus values corresponding to regions outside of condensates (i.e. the mica surface) were excluded from analysis.

### 4.8. Fluorescence Correlation Spectroscopy (FCS)

For FCS analysis, slides were prepared in the same manner as described above for the confocal imaging experiment. Using the same Leica AOBS SP8 FALCON confocal microscope with a 63x/1.2 water objective, FCS measurements were taken five consecutive times at three separate points within condensates (center, intermediate, edge), and data was normalized by applying the structural parameter and effective volume values generated by calibrating the system using known diffusion coefficients and free fluorophores (Alexa Fluor 488: 435 μm2/s; Alexa Fluor 647: 440 μm2/s). Leica Application Suite X v3.5 (LASX, Leica Microsystems, Wetzlar, Germany) software with built-in FCS functionality was used to collect and analyze data.

### 4.9. Quantitative Image and Statistical Analyses

Image analysis was performed in Leica Application Suite X v3.5 (LASX, Leica Microsystems, Wetzlar, Germany), Imaris Image Analysis Software v10.2.0 (Oxford Instruments, Abingdon, United Kingdom), or Bruker BioAFM Data Processing Software v8.0 (Bruker, Billerica, MA, USA) as indicated. Huygens Deconvolution software v24.10 (Scientific Volume Imaging, Hilversum, Netherlands) was utilized for deconvolution. Data were analyzed and plotted using Microsoft Excel (Microsoft, Redmond, WA, USA) and GraphPad Prism v10.2.2 (GraphPad Software, Boston, MA, USA).

## Supplementary Materials

Video S1: RSV Gag Alone Confocal Morphology.mp4;

Video S2: RSV Gag + gRNA Ψ DIS Confocal Morphology.mp4;

Video S3: RSV Gag + gRNA Ψ ΔDIS Confocal Morphology.mp4

## Acknowledgments

The authors would like to thank John Flanagan, PhD (Department of Molecular and Precision Medicine, Penn State College of Medicine), for purifying the RSV Gag protein; Malgorzata Sudol, MS (Department of Medicine, Penn State College of Medicine), for technical assistance; Lichong Xu, PhD (Department of Surgery, Penn State College of Medicine) for AFM training and assistance; Vonn Walter, PhD (Department of Public Health Sciences, Penn State College of Medicine) and the Biostatistics/Epidemiology/Research Design (BERD) Core of the Penn State Clinical and Translational Science Institute (CTSI) for assistance with statistical analysis; and Rebecca J. Kaddis Maldonado, PhD (Departments of Medicine and Molecular and Precision Medicine, Penn State College of Medicine) and Alecia Achimovich, PhD (Department of Medicine, Penn State College of Medicine) for their constructive feedback. The Advanced Light Microscopy Core (RRID:SCR_022526) and Atomic Force Microscopy Core (RRID:SCR_025075) services and instruments used in this project were funded, in part, by The Pennsylvania State University College of Medicine via the Office of the Vice Dean of Research and Graduate Students and the Pennsylvania Department of Health using Tobacco Settlement Funds (CURE, https://www.pa.gov/agencies/health/research/research/cure.html). The content is solely the responsibility of the authors and does not necessarily represent the official views of the University or College of Medicine. The Pennsylvania Department of Health specifically disclaims responsibility for any analyses, interpretations or conclusions. We acknowledge support for the Atomic Force Microscopy Core through an NIH S10 award (1S10OD030279-01A1).

